# Perinatal Semaglutide Treatment Improves Maternal Health and Mitigates Offspring Metabolic Dysfunction in a Mouse Model of Maternal Obesity

**DOI:** 10.64898/2026.06.07.730723

**Authors:** Ananthi Rajamoorthi, Taylor Hollingsworth, Yuxia Guan, Sara E. Pinney, Rebecca A. Simmons

## Abstract

Early-life exposures during critical periods of development significantly impact lifelong metabolic risk and likely contribute to the rising rates of obesity, type 2 diabetes, and metabolic dysfunction–associated steatotic liver disease (MASLD) in children. Here, we evaluated the safety and metabolic effects of semaglutide, a GLP-1 receptor agonist (GLP-1 RA), administered from preconception through lactation in dams fed a high-fat diet (HFD) or standard diet, and assessed metabolic outcomes in dams and their offspring. Offspring were weaned to a standard diet. We found that semaglutide improved body composition and glucose metabolism in HFD-fed dams during pregnancy. These maternal changes persisted 10 weeks after weaning despite discontinuation of semaglutide treatment. HFD exposure impaired glucose homeostasis and promoted hepatic steatosis in offspring at 18 weeks. These effects were ameliorated by maternal semaglutide treatment. Importantly, metabolic improvements in dams and offspring occurred without adverse effects on conception rate or fetal viability. These findings suggest that GLP-1 RA during the perinatal period can improve maternal and offspring metabolic health in a mouse model of obesity and support further investigation of GLP-1–based therapies to mitigate maternal metabolic dysfunction and improve metabolic risk in children.

**ARTICLE HIGHLIGHTS:**

• Rates of obesity, type 2 diabetes, and fatty liver disease are rising in children, in part due to maternal obesity and insulin resistance that program offspring metabolic risk during the perinatal period.
• We asked whether the GLP-1 receptor agonist (GLP-1 RA), semaglutide, administered during critical developmental windows could prevent adverse outcomes in offspring using a diet-induced mouse model of maternal obesity.
• Semaglutide, given to dams from preconception through lactation, improved maternal metabolism and ameliorated metabolic dysfunction in offspring caused by maternal high-fat diet.
• These findings highlight a potential role for perinatal GLP-1 receptor agonism to improve maternal metabolic health and reduce metabolic risk in offspring.

## INTRODUCTION

Metabolic syndrome is defined by co-occurring metabolic conditions, including obesity, insulin resistance, dyslipidemia, and hypertension, and affects almost 67 million children worldwide (1, 2). The World Health Organization recently reported that the pediatric obesity rate has quadrupled since 1990, while youth onset Type 2 diabetes incidence in the US has increased over 5-fold from 2001 to 2017 (3, 4). Pediatric metabolic dysfunction–associated steatotic liver disease (MASLD), most prevalent among children with overweight or obesity, affects 10% of US children and represents the leading cause of chronic liver disease in youth worldwide (5, 6). For these reasons, many consider pediatric metabolic disease as a public health emergency (7).

A subset of children develop metabolic dysfunction–associated steatohepatitis (MASH), a progressive form of steatotic liver disease characterized by inflammation and fibrosis, which is more severe and associated with earlier fibrosis compared with adult-onset disease (5, 8, 9). A similar pattern of earlier onset of severe disease is seen in youth onset T2DM, suggesting that childhood metabolic dysfunction poses a unique public health challenge, with distinct pathophysiology from adult metabolic disease (3, 10). This highlights a need for both targeted intervention and more focused prevention strategies, which are currently lacking. There is strong evidence that the accelerated progression and increased severity of early onset metabolic dysfunction is in part due to intrauterine and early postnatal factors that affect the developing cells in the pancreas, liver, and adipose tissue, thus programming individuals to develop early onset metabolic disease (11–16). This concept, known as the developmental origins of health and disease (DOHaD), suggests that critical periods of development are particularly amenable to disruption by maternal factors (12, 16, 17). In both preclinical and large population based clinical studies, maternal factors have been shown to independently program offspring metabolic risk including maternal obesity, glucose, insulin, insulin resistance, and adiponectin (18–26). Clinically, rising rates of maternal obesity, gestational diabetes mellitus, and even subclinical metabolic dysfunction characterized by higher maternal glucose and insulin levels during pregnancy independent of BMI, are associated with pregnancy complications and offspring metabolic dysfunction (19, 27, 28).

Our lab previously showed that early postnatal treatment of growth restricted pups with GLP-1 RA (exendin-4) normalized β-cell development and insulin secretion and prevented hepatic insulin resistance in offspring exposed to uteroplacental insufficiency (29, 30). Building on this foundation, we tested the hypothesis that preconception semaglutide treatment of diet-induced obese C57BL/6 mice improves maternal and offspring metabolic homeostasis. We show that maternal semaglutide at clinically relevant doses improves maternal body composition and glycemic control during gestation and postpartum, and prevents offspring accelerated postnatal weight gain, glucose intolerance, and hepatic steatosis in adulthood. These data suggest that GLP-1 RA treatment represents a promising strategy to simultaneously improve maternal metabolic health while mitigating the transmission of obesity and metabolic disease risk to offspring.

## RESEARCH DESIGN AND METHODS

### Chemicals

Vehicle and GLP-1 receptor agonist (GLP-1 RA), semaglutide, were provided by Novo Nordisk and stored at -80°C until thawed immediately prior to use. For glucose tolerance testing and pyruvate challenge test, glucose and sodium pyruvate were purchased from Sigma (St. Louis, MO). For insulin tolerance testing, regular insulin was used (Humulin-R, Eli Lilly, Indianapolis, IN).

### Experimental Animals

C57BL/6 mice were obtained from The Jackson Laboratory (JAX) at 6 weeks of age and acclimated to the animal facility for 2 weeks prior to experiments. All animals were maintained in a 12 h/12 h light/dark cycle with ad libitum access to food and water. Eight-week-old C57BL/6 female mice were fed either a chow diet or a high-fat diet (HFD; TD.06414, Inotiv, USA) for 10 weeks (Figure 1A). Mice received vehicle or a clinically relevant dose of semaglutide (3 nmol/kg) via subcutaneous injection for 5 days. On the fifth day of treatment, female mice were paired with chow-fed males overnight, and pregnancy was confirmed by vaginal plug (embryonic day [E] 0.5). During gestation, dosing of vehicle or semaglutide was reduced to 1 nmol/kg. This dosing regimen was selected based on dosing guidance provided by scientists at Novo Nordisk and prior published preclinical studies (31–33). Glucose tolerance (GTT) and insulin tolerance tests (ITT) were performed in pregnant dams at E14.5–E16.5. The first cohort of pregnant females (*n* = 5 chow, *n* = 8 HFD) was sacrificed at E17.5 and maternal plasma and tissues (placenta, liver, and gonadal white adipose tissue) were collected. In a second, independent cohort of dams (*n* = 5-7 chow, *n* = 7-8 HFD), treatment with vehicle or semaglutide was continued throughout lactation at 1 nmol/kg, with dosing withheld during the first 3 days postpartum to minimize cannibalism.

**Figure 1.**
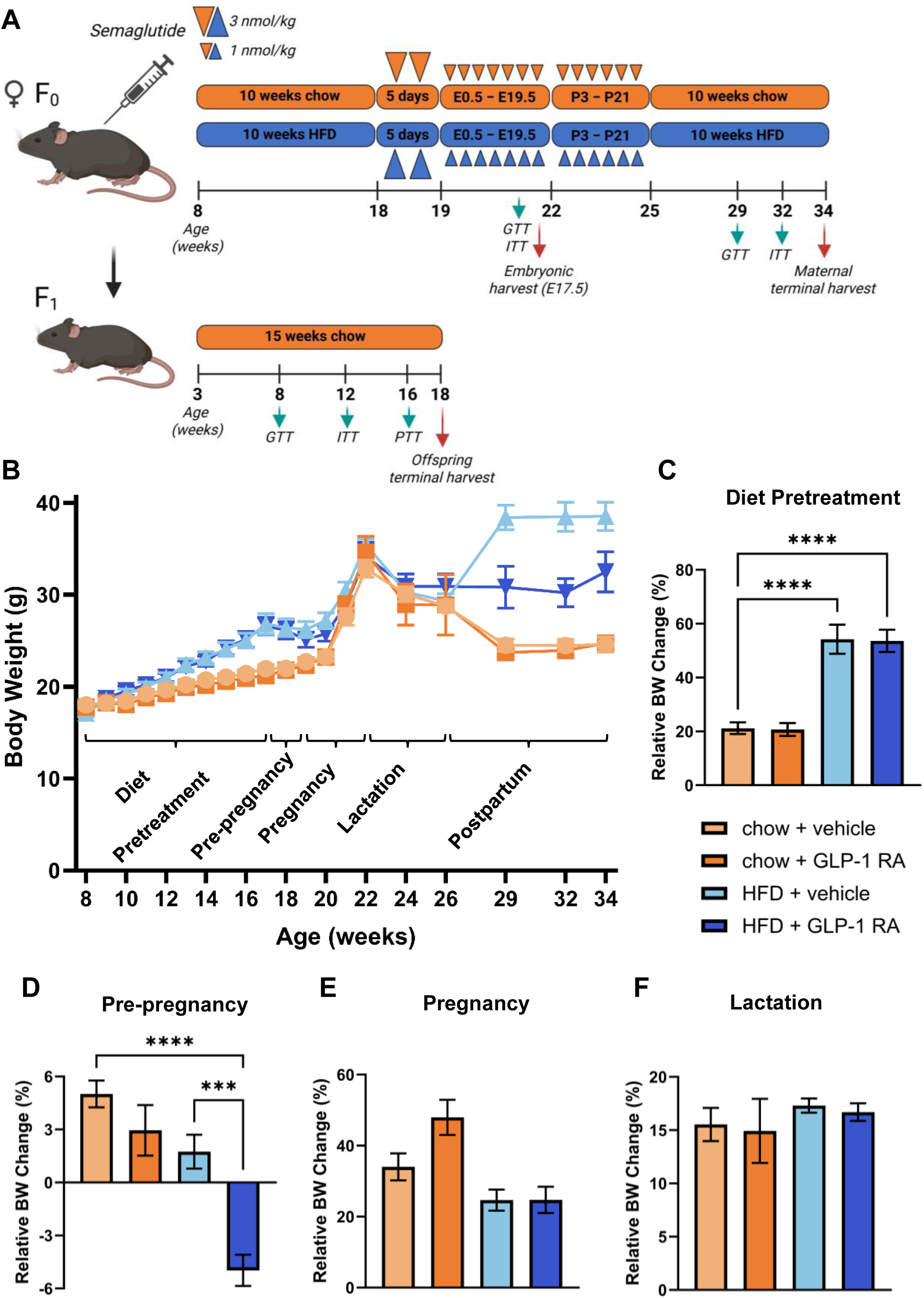
Experimental design and body weight changes in dams. (A) Experimental design. Eight-week-old female C57BL/6 mice were maintained on chow or high-fat diet (HFD) for 10 weeks (“Diet Pretreatment”). Semaglutide was administered at 3 nmol/kg before conception (“Pre-pregnancy”) and reduced to 1 nmol/kg during pregnancy (“Pregnancy”) and lactation (“Lactation”). Treatment was discontinued after weaning (“Postpartum”). A subset of dams was sacrificed at embryonic day (E)17.5 for maternal tissue analyses, while a second cohort underwent longitudinal maternal and offspring metabolic phenotyping. F1 offspring were weaned onto chow diet and studied through 18 weeks of age. (B) Body weight trajectory of dams from pretreatment through the postpartum period. (C–F) Relative body weight change during the diet pretreatment (C), prepregnancy (D), pregnancy (E), and lactation (F) phases of the study. Data are presented as mean ± SEM (*n* = 10–12 chow, *n* = 15–16 HFD). \*\*\**P* ≤ 0.001, \*\*\*\**P* ≤ 0.0001. GTT, glucose tolerance test; ITT, insulin tolerance test; PTT, pyruvate tolerance test.

Following weaning, maternal treatment was discontinued. GTT and ITT were performed in dams at 5 and 8 weeks postweaning. Dams were sacrificed at 34 weeks of age, 10 weeks after weaning. Dams from both cohorts were sacrificed during the light cycle in the fed state. Body weight was measured in dams throughout the study, and food intake was quantified at specified time points (preconception day 5 and gestational day E10.5) as the difference between food provided and food remaining over a 24-hour period.

F1 offspring from the second cohort of dams across all four maternal exposure groups—chow + vehicle, chow + GLP-1 RA, HFD + vehicle, and HFD + GLP-1 RA—were weaned onto chow at 3 weeks of age. GTT, ITT, and pyruvate tolerance tests (PTT) were performed at 8, 12, and 16 weeks of age, respectively. Offspring were sacrificed at 18 weeks of age following a 6-hour fast during the light cycle. Offspring plasma and liver tissue were collected. Body weight was measured in offspring throughout the study. Body composition and bone density were measured in dams and offspring via EchoMRI and DEXA PIXImus on the days of sacrifice at the University of Pennsylvania Rodent Metabolic Phenotyping Core. Diet composition is as follows: chow diet (5LOD), 2.86 kcal/g (28.9% protein, 57.5% carbohydrate, 13.6% fat); HFD (TD.06414), 5.1 kcal/g (18.3% protein, 21.4% carbohydrate, 60.3% fat), with fat content derived primarily from lard and soybean oil. Animal studies were conducted in conformity with the Public Health Service policy on humane care and use of laboratory animals and approved by the IACUC at the Children’s Hospital of Philadelphia.

### Glucose, insulin, and pyruvate challenges

GTTs were performed in dams (P0) fasted for 4 hours and offspring (F1) fasted for 6 hours during the dark cycle and then underwent intraperitoneal (ip) injection with 2 g/kg glucose. ITTs were performed in dams and offspring fasted for 2 hours during the light cycle then injected ip with Humulin-R at 0.75 U/kg in dams (P0) or 0.70 U/kg in offspring (F1). Pyruvate tolerance tests (PTTs) were performed in male offspring (F1) fasted for 16 hours overnight and then injected ip with 2 g/kg sodium pyruvate. In all studies, glycemia was monitored at the indicated time points from tail bleeds using a One Touch Ultra Blue (LifeScan U.S. LLC). Absolute area under the curve (AUC) was calculated for GTTs. For ITTs, blood glucose was normalized to baseline (0 min) and expressed as percentage of baseline values prior to calculation of relative AUC to assess systemic insulin sensitivity independently of differences in fasting glucose.

### Histology

Samples of liver, gonadal white adipose tissue, and placenta were fixed in 4% paraformaldehyde, postfixed in 75% ethanol, and embedded in paraffin blocks. Sections (4 μm) were processed for hematoxylin and eosin (H&E) staining using standard techniques. Frozen liver and placenta sections were used for Oil-Red-O (ORO) lipid staining. ImageJ was used for quantitative image analysis, including assessment of ORO staining in liver and placenta sections, measurement of adipocyte surface area, and evaluation of placental morphology parameters such as junctional zone (JZ) thickness and labyrinth (Lb)-to-JZ ratio (34).

### Lipid analysis

Hepatic triglyceride content was quantified using a commercial enzymatic assay kit (Cayman Chemical, Ann Arbor, MI). Briefly, liver tissue (175–200 mg) was homogenized in NP-40–based lysis buffer with protease inhibitor, and triglycerides were measured enzymatically following the manufacturer’s instructions. In this assay, triglycerides are hydrolyzed to glycerol and free fatty acids, and glycerol is subsequently quantified through an enzymatic colorimetric reaction, with absorbance measured on a microplate reader and compared against a triglyceride standard curve. Triglyceride content was normalized to liver tissue weight.

### Plasma analysis

Plasma insulin was measured using an ELISA kit (Proteintech KE10089, Rosemont, IL).

### RNA analysis

RNA was isolated from liver using Qiagen RNeasy Kit and analyzed by real-time quantitative PCR using Fast SYBR Green (Life Technologies, Carlsbad, CA) and a QuantStudio 7 Flex Real-Time PCR System (Applied Biosystems, Thermo Fisher Scientific, Waltham, MA). Values were normalized to 36b4, and relative expression calculated using the ΔΔCT method. Primer sets are shown in supplemental Table S1.

### Statistics

Data are presented as mean ± SEM. Statistical analyses for continuous variables were performed using one-way ANOVA with maternal exposure group (chow + vehicle, chow + GLP-1 RA, HFD + vehicle, and HFD + GLP-1 RA) as the independent variable, followed by Tukey’s multiple comparisons test. Categorical variables were analyzed using Fisher’s exact test, as appropriate. A P value < 0.05 was considered statistically significant. No a priori sample size calculation was performed; sample sizes were based on prior studies and experimental feasibility.

## RESULTS

### Maternal semaglutide improved adiposity and glycemic control in HFD-fed mice during gestation

To determine whether maternal semaglutide altered maternal body weight, body composition, and food intake, dams were treated with semaglutide before and during gestation and maternal parameters were measured throughout study period (Figure 1A). Absolute body weight trajectory shows notable differences in body weight among groups (Figure 1B). Female mice fed a HFD for 10 weeks, as expected, gained 55% relative body weight (Figure 1C). Semaglutide treatment in HFD-fed animals pre-pregnancy resulted in modest weight loss of 5% with 5 days of treatment at 3 nmol/kg dosing (Figure 1D). Treatment at lower dosing (1 nmol/kg) during gestation (Figure 1E) and lactation (Figure 1F) did not result in further weight loss during these study periods in either chow- or HFD-fed animals.

Food intake mirrored changes in body weight. Specifically, while pre-pregnancy semaglutide treatment resulted in a 34% reduction in food intake on day 5 of treatment (Supplementary Figure 1A), no significant differences in food intake were noted at E10.5 (Supplementary Figure 1B).

Changes in fat mass in HFD- fed female mice paralleled body weight trajectory, without changes in lean mass during gestation. Specifically, there was a trend for reduced fat mass, adjusted for litter size, in HFD-fed animals treated with semaglutide (Figure 2A). Interestingly, chow-fed female mice treated with semaglutide had a higher lean mass during gestation (Figure 2A). Treatment significantly reduced relative gonadal white adipose tissue (gWAT) weight in HFD-fed animals (Figure 2B), as well as mean adipocyte surface area (Figures 2C and 2D), despite the absence of changes in relative body weight during gestation (Figure 1E). In addition, maternal semaglutide treatment resulted in a trend for improved glucose tolerance in chow-fed animals, with a significant difference noted with treatment in the HFD group (Figure 2E). Insulin tolerance test did not show statistically significance differences when taking into consideration differences in baseline glycemia with treatment in HFD-fed animals (Figure 2F).

**Figure 2.**
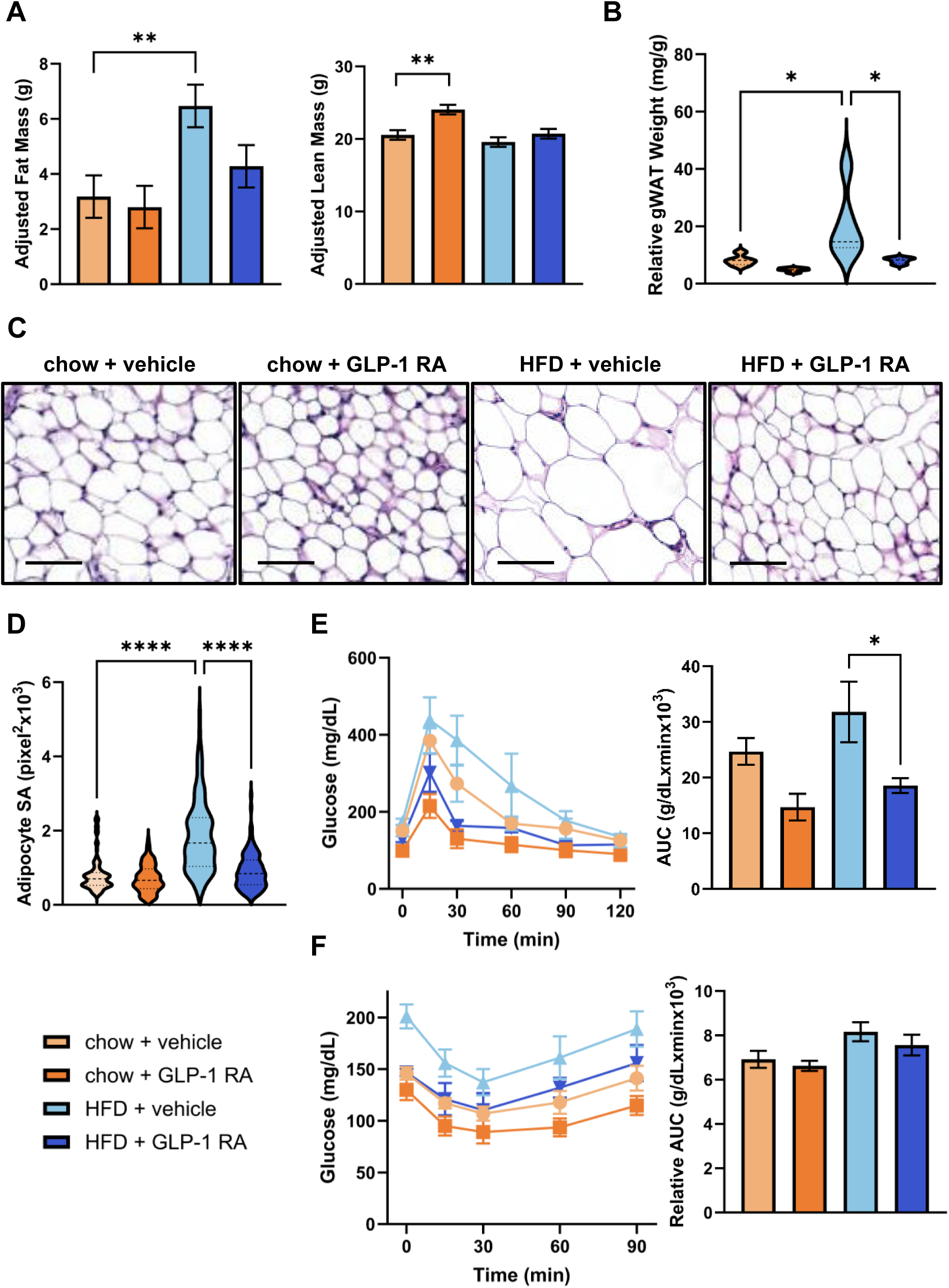
Maternal adiposity and glucose homeostasis during pregnancy. (A) Adjusted fat mass and lean mass measured by MRI in pregnant dams at embryonic day (E)17.5, accounting for variations in litter size. (B) Relative gonadal white adipose tissue (gWAT) weight at E17.5. (C) Representative hematoxylin and eosin (H&E)-stained gonadal white adipose tissue sections from pregnant dams at E17.5. Scale bar, 100 μm. (D) Quantification of mean adipocyte surface area (SA) from gonadal WAT sections. (E) Glucose tolerance test (GTT) and (F) insulin tolerance test (ITT) performed in pregnant dams at E14.5–E15.5, with corresponding area under the curve (AUC) for GTT and relative AUC for ITT. Data are presented as mean ± SEM (*n* = 5 chow, *n* = 8 HFD). *P ≤ 0.05, **P ≤ 0.01, ****P ≤ 0.0001.

### Maternal semaglutide reduced hepatic steatosis in HFD-fed mice during gestation

To assess whether maternal semaglutide altered hepatic lipid content in dams at E17.5, hematoxylin and eosin (H&E) and Oil-Red-O (ORO) staining were performed in liver sections, which showed a significant increase in neutral lipid staining with HFD that was reduced with semaglutide treatment (Figures 3A and 3B). Consistent with diet-induced steatosis, dams fed a HFD had significantly elevated triacylgycerol (TAG) content that was reduced with semaglutide treatment (Figure 3C). Relative liver weight was reduced with HFD (Figure 3D), despite similar absolute liver weights among groups (data not shown), suggesting this difference was driven primarily by changes in overall body weight. Greater liver lipid content with HFD was accompanied by increased mRNA expression of lipid droplet-associated markers *Cidea* and *Fsp27*, as well as the pro-inflammatory cytokine *Il1b* and profibrotic marker *Timp1*. Semaglutide treatment during pregnancy and gestation in HFD-fed animals resulted in decreased expression of the lipid transport marker *Cd36*, as well as *Cidea* and *Fsp27*, and inflammatory and fibrotic markers *Il1b* and *Timp1* (Figure 3E). Interestingly, although semaglutide treatment in chow-fed mice reduced hepatic TAG content, no overt histologic differences in hepatic lipid accumulation or ORO staining were observed compared with chow + vehicle controls, likely reflecting the relatively low baseline hepatic lipid burden in chow-fed animals. This reduction in hepatic TAG content in chow-fed mice treated with semaglutide was accompanied by reduced expression across lipogenic, oxidative, and lipid droplet-associated transcripts, as well as proinflammatory cytokine *Tnfa* and *Timp1*.

**Figure 3.**
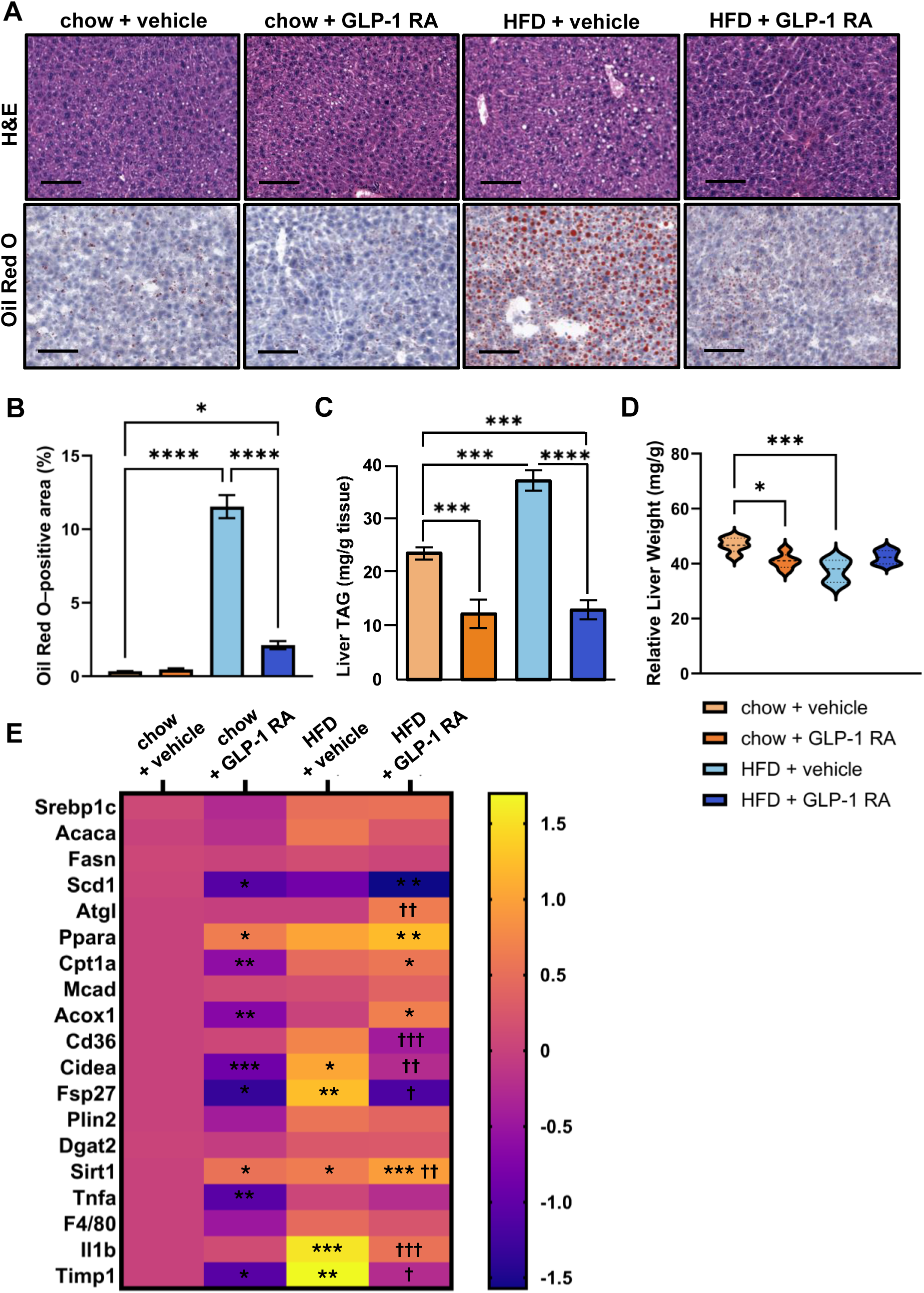
Maternal hepatic lipid accumulation and gene expression during pregnancy. (A) Representative hematoxylin and eosin (H&E, top) and Oil Red O (ORO, bottom) staining of maternal liver sections at embryonic day (E)17.5. Scale bar, 100 μm. (B) Quantification of ORO-positive area in liver sections. (C) Hepatic triglyceride (TAG) content. (D) Relative liver weight at E17.5. (E) Hepatic gene expression profiling of lipogenic, oxidative, lipid storage and transport, inflammation, and fibrosis markers assessed by bulk quantitative PCR (qPCR), presented as log2 fold change in a heatmap. Data are presented as mean ± SEM (*n* = 5 chow, *n* = 8 HFD). *P ≤ 0.05, **P ≤ 0.01, ***P ≤ 0.001, ****P ≤ 0.0001 vs chow + vehicle. ^†^P ≤ 0.05, ^††^P ≤ 0.01, ^†††^P ≤ 0.001 vs HFD + vehicle.

### Maternal semaglutide did not impair conception or fetal viability

To determine whether maternal semaglutide affected pregnancy outcomes in chow- and HFD-fed animals, conception rate, fetal viability, and fetal and placental weights were assessed at E17.5. No statistically significant differences in conception rate were observed among the four groups (Supplementary Figure 2A). While there were no significant differences in total fetuses or resorptions per litter, semaglutide treatment in chow-fed animals resulted in an increase in viable fetuses per dam compared to chow + vehicle group, although no differences in fetal viability were noted (Supplementary Figures 2B and 2C). Neither diet nor treatment significantly affected fetal weight. However, semaglutide treatment in both chow- and HFD-exposed animals reduced placental weight, which in the chow group, corresponded to an increase in fetal to placental weight ratio (Supplementary Figure 2D).

To assess effects of diet and treatment on placental architecture, junctional zone (JZ) thickness and labyrinth to junctional zone (Lb:JZ) ratio were measured in placental cross sections (Supplementary Figure 2E). HFD resulted in an increase in JZ thickness.

Chow + semaglutide group had a greater Lb:JZ ratio compared to chow + vehicle, with no differences seen with semaglutide treatment in HFD-fed group (Supplementary Figure 2F). ORO staining of placental cross sections revealed a significant increase in neutral lipid content within maternal decidua in HFD + vehicle group compared to chow control group, that was mitigated by semaglutide treatment (Figure 2G).

### Semaglutide during gestation and lactation reduced adiposity and improved glycemic control in dams postpartum

To determine whether exposure to semaglutide preconception, and during gestation and lactation, modified metabolic parameters in female mice (F0) after discontinuing treatment upon weaning offspring in the postpartum period, body weight, body composition, and glucose and insulin tolerance were measured. After discontinuing treatment, primiparous female mice (F0) on chow diet lost body weight as expected, and animals on HFD gained weight during this period compared to chow controls, which was mitigated by prior semaglutide exposure (Figure 4A). Body composition analysis revealed notable differences in fat mass 10 weeks after discontinuing treatment. Specifically, female mice previously treated with semaglutide before and during gestation and lactation had a 32% lower absolute fat mass compared with HFD-fed animals previously treated with vehicle (Figure 4B). These changes occurred in the absence of changes in lean mass or bone density (Figures 4B and 4C). Glucose and insulin tolerance tests performed 4 and 8 weeks postpartum, respectively, revealed notable differences in glucose handling, with HFD + vehicle group displaying significant impairment in glycemic control (Figure 4D). Despite discontinuation of treatment for 4 weeks post-weaning, HFD + GLP-1 RA group showed significant improvement in glycemia across time points, compared to HFD + vehicle group (Figure 4D). These differences among groups were also observed with insulin tolerance test measured 8 weeks after parturition, suggesting persistent alterations in maternal insulin sensitivity postpartum despite discontinuation of semaglutide (Figure 4E).

**Figure 4.**
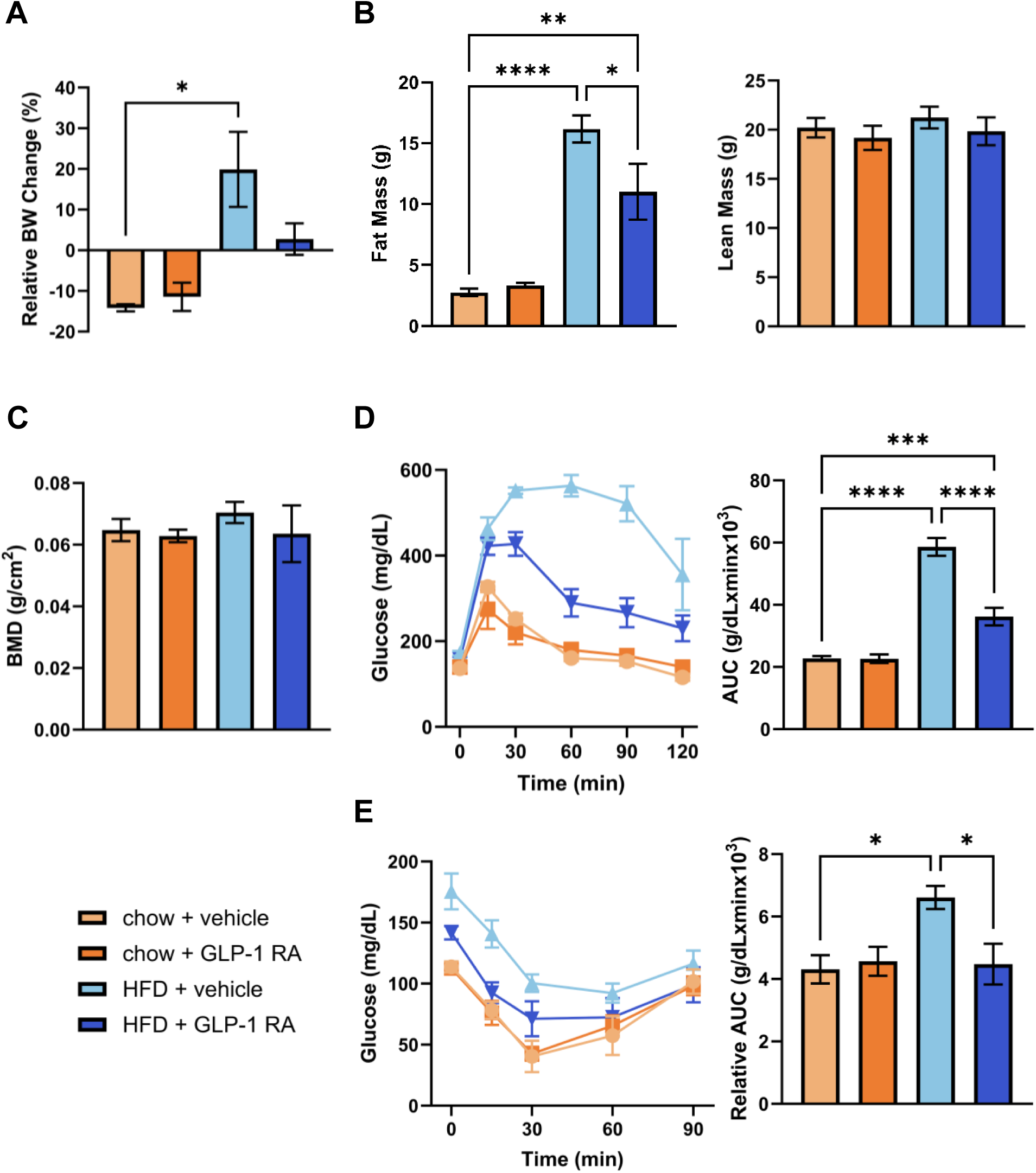
Postpartum metabolic phenotype in primiparous dams. (A) Relative body weight change during the postpartum period. (B) Absolute fat mass and lean mass measured by MRI and (C) Bone mineral density (BMD) measured by DEXA, in adult primiparous female mice assessed 10 weeks after parturition. (D) Glucose tolerance test (GTT) performed 4 weeks postpartum, and (E) Insulin tolerance test (ITT) performed 8 weeks postpartum, with corresponding area under the curve (AUC) for GTT and relative AUC for ITT. Data are presented as mean ± SEM (*n* = 5–7 chow, *n* = 7–8 HFD). *P ≤ 0.05, **P ≤ 0.01, ***P ≤ 0.001, ****P ≤ 0.0001.

### Maternal semaglutide prevented postnatal accelerated weight gain and reduced adiposity in male offspring exposed to HFD during gestation and lactation

To assess the effects of maternal semaglutide treatment on offspring (F1) outcomes, body weight was measured longitudinally until 18 weeks of age. Offspring, including both male and female pups, exposed to HFD during gestation and lactation showed an accelerated pattern of weight gain in the immediate postnatal period from P7 through P21, which was prevented by maternal semaglutide treatment (Figures 5A and 5B). While there were no significant changes in absolute body weight beyond the postnatal period through 18 weeks of age (Figure 5C), HFD + vehicle male offspring had greater fat mass compared to chow + vehicle group (Figure 5D). Importantly, HFD + semaglutide male offspring had near normal fat mass similar to chow + vehicle offspring, and semaglutide exposure in both the chow and HFD groups did not impact lean mass or BMD in male offspring (Figures 5D and 5E). Reductions in fat mass with maternal semaglutide in HFD-exposed male offspring were further reflected in gross reductions in relative epididymal white adipose tissue (eWAT) weight at 18 weeks of age (Figure 5F), as well as significant reductions in mean adipocyte surface area compared with HFD + vehicle male offspring (Figures 5G and 5H). Notable differences in adipocyte surface area (SA) distribution were observed among groups (Figure 5I). Male offspring exposed to HFD during gestation and lactation exhibited a reduced frequency of smaller adipocytes and an increased frequency of larger adipocytes compared with controls. These shifts in adipocyte size distribution were attenuated in semaglutide-exposed animals.

**Figure 5.**
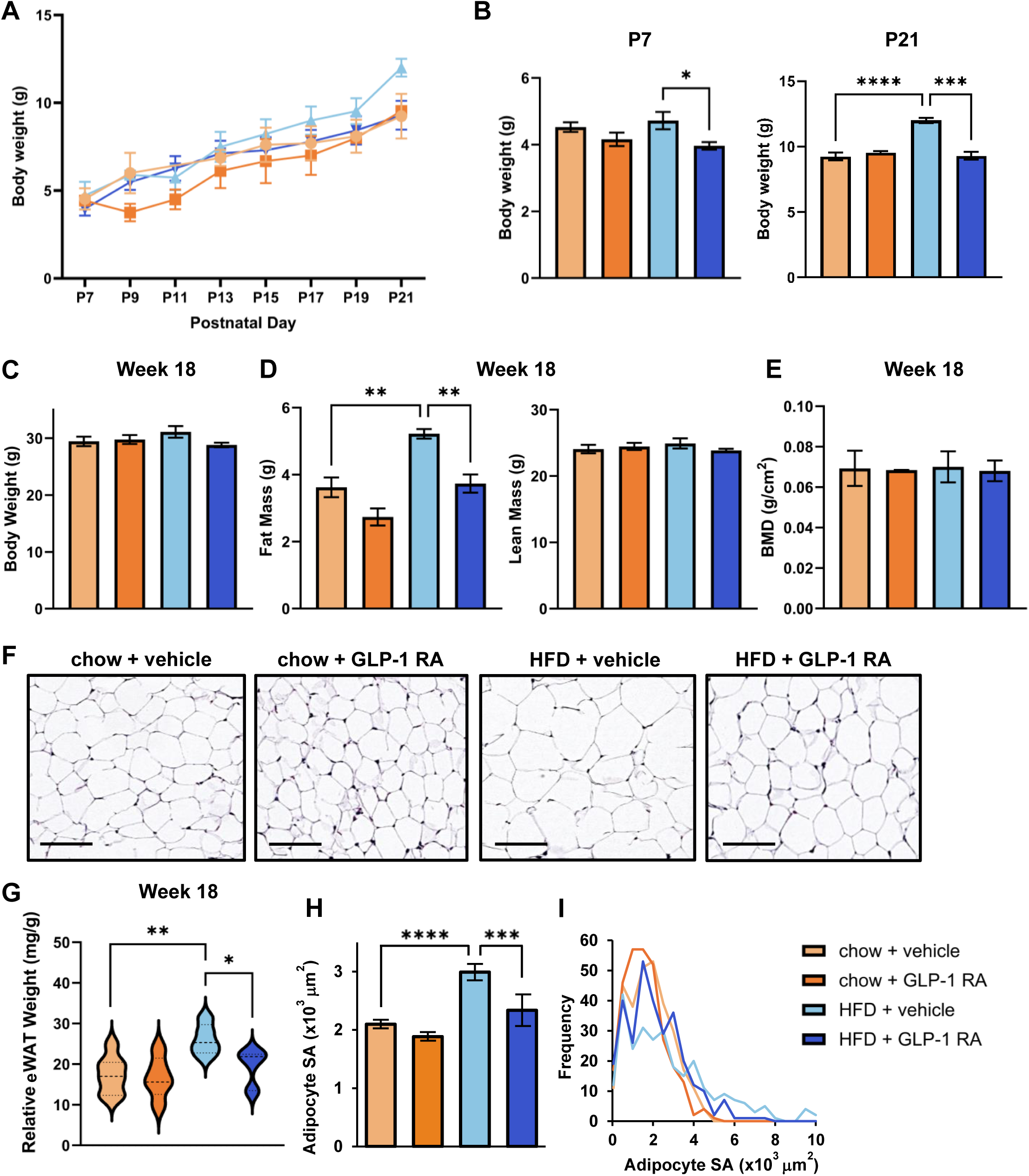
Postnatal growth, body composition, and adipose tissue morphology in F1 offspring. (A) Postnatal body weight trajectory in F1 offspring. (B) Average body weight at postnatal day (P)7 and P21. Male and female offspring are included in (A) and (B). (C–I) Male offspring analyses at 18 weeks of age, including (C) body weight, (D) absolute fat and lean mass measured by MRI, (E) bone mineral density (BMD) measured by DEXA, (F) relative epididymal white adipose tissue (eWAT) weight, (G) representative hematoxylin and eosin (H&E)-stained eWAT sections (scale bar, 100 μm), (H) mean adipocyte surface area (SA), and (I) distribution of adipocyte surface area shown as frequency (cell count) across size bins. Data are presented as mean ± SEM (A, B: *n* = 5–7 litters; C–I: *n* = 6–9 male offspring). *P ≤ 0.05, **P ≤ 0.01, ***P ≤ 0.001, ****P ≤ 0.0001.

Female offspring exposed to maternal HFD did not exhibit significant differences in body weight, adiposity, or glucose tolerance compared with controls. A modest impairment in insulin tolerance was observed in female HFD-exposed offspring (Supplementary Fig. 3). Given the more pronounced metabolic phenotype observed in male offspring, subsequent analyses were focused on male offspring to further evaluate metabolic effects of maternal GLP-1 RA exposure during pregnancy and lactation.

### Maternal semaglutide improved glycemic control in adult male offspring exposed to HFD during gestation and lactation

To assess the effects of diet and treatment on fasting metabolic parameters and glucose handling in adult offspring, fasting glucose and insulin levels were first measured, followed by glucose, insulin, and pyruvate tolerance tests in male offspring at 8, 12, and 16 weeks of age, respectively. Offspring exposed to HFD + vehicle exhibited a trend toward increased fasting glucose compared with chow + vehicle controls (Table 1, 163.3 ± 9.6 vs. 145.2 ± 8.5 mg/dL, P = 0.187). Fasting insulin was significantly elevated in HFD + vehicle offspring compared with chow + vehicle controls (5.6 ± 0.7 vs. 3.81 ± 0.69 ng/mL, P = 0.003), whereas HFD + semaglutide offspring exhibited significantly lower fasting glucose (137.6 ± 7.9 mg/dL, P = 0.048) and insulin (3.49 ± 0.46, P = 0.019) compared with HFD + vehicle controls. Semaglutide exposure also showed a trend toward lower fasting glucose and insulin in chow-fed offspring. Consistent with improved fasting metabolic parameters, male offspring exposed to HFD + semaglutide showed improved glucose, insulin, and pyruvate tolerance compared with HFD + vehicle (Figures 6A–6C).

**Figure 6.**
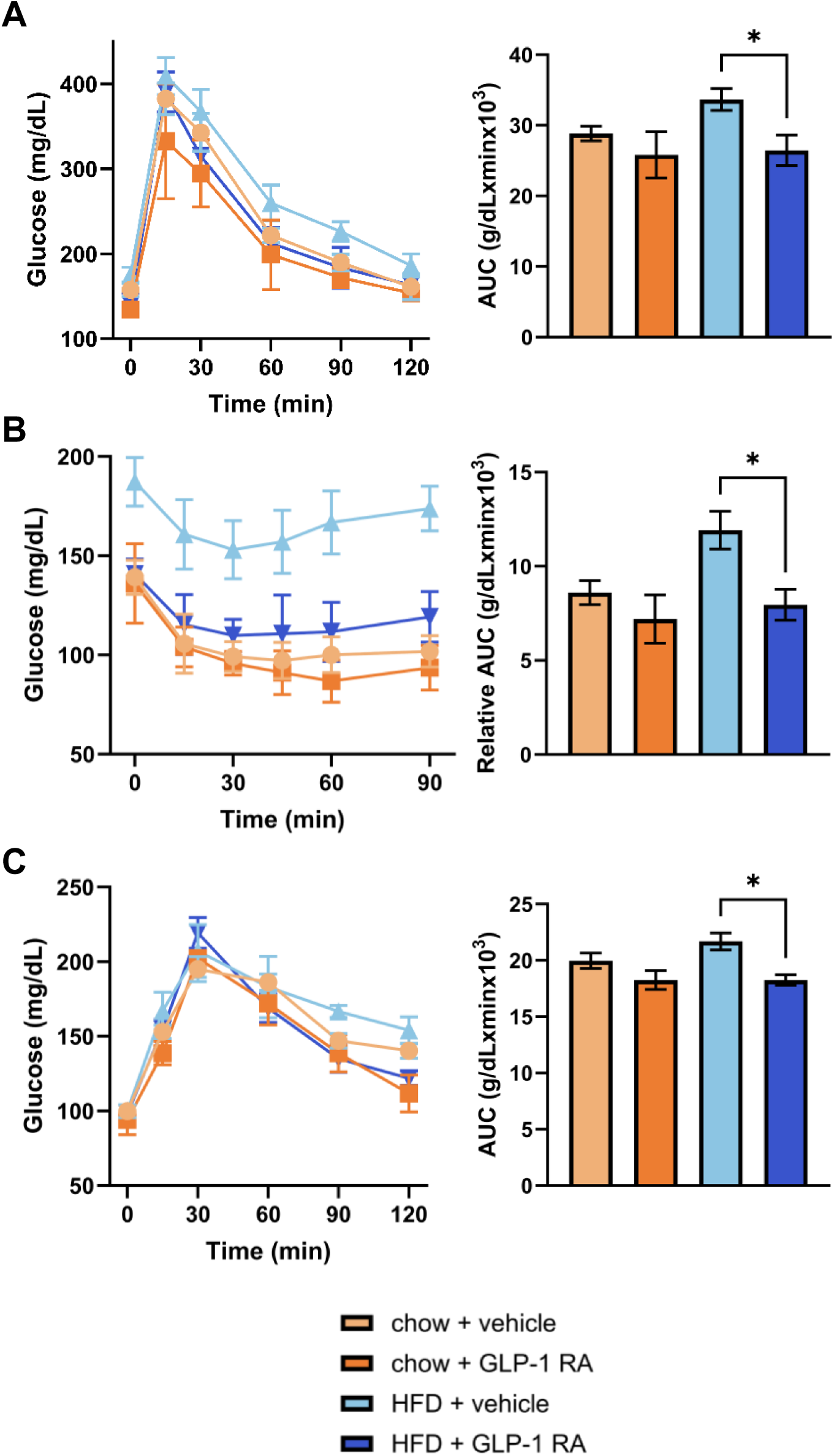
Glucose, insulin, and pyruvate tolerance in male F1 offspring. (A) Glucose tolerance test (GTT) at 8 weeks, (B) insulin tolerance test (ITT) at 12 weeks, and (C) pyruvate tolerance test (PTT) at 16 weeks in male F1 offspring. Area under the curve (AUC) was calculated for GTT and PTT, and relative AUC was calculated for ITT. Data are presented as mean ± SEM (*n* = 6–9 male offspring). *P ≤ 0.05.

**Table 1.** Serum metabolic markers in male F1 offspring at 18 weeks of age. Fasting glucose and fasting insulin were measured after a 6-hour fast during the light cycle. Data are presented as mean ± SEM. **P ≤ 0.01 (vs chow + vehicle). ‡P ≤ 0.05 (vs HFD + vehicle).

|  | chow + vehicle | chow + GLP-1 RA | HFD + vehicle | HFD + GLP-1 RA |
| --- | --- | --- | --- | --- |
| **Fasting glucose (mg/dL)** | 145.2 ± 8.5 | 119.0 ± 5.0 | 163.3 ± 9.6 | 137.6 ± 7.9‡ |
| **Fasting insulin (ng/mL)** | 3.81 ± 0.69 | 2.49 ± 0.80 | 5.6 ± 0.70** | 3.49 ± 0.46‡ |

### Maternal semaglutide reduced steatosis in adult male offspring exposed to HFD during gestation and lactation

To assess effects of maternal diet and treatment on hepatic lipid content in adult offspring at 18 weeks, H&E and ORO staining were performed in liver sections from adult male offspring, which showed an increase in hepatic neutral lipid staining and TAG content with HFD compared to chow + vehicle controls, that was reduced in male offspring exposed to semaglutide (Figures 7A and 7B). Consistent with diet-induced steatosis, dams fed a HFD showed a trend for elevated TAG content that was reduced with semaglutide treatment (Figure 7C). No differences in relative liver weight were observed among groups (Figure 7D). Male offspring livers exposed to HFD during gestation and lactation had increased expression of *Cidea* and lipid transporter *Cd36*, as well as increased expression of *Tnfa*, compared to chow + vehicle controls. These transcripts were reduced in livers from male offspring exposed to both HFD and semaglutide during gestation and lactation. HFD + semaglutide group also showed increased mRNA expression of lipolytic and oxidative markers including *Atgl*, *Ppara*, and *Mcad* as well as a reduction in lipogenic transcript *Srebp1c*, compared to chow + vehicle group.

**Figure 7.**
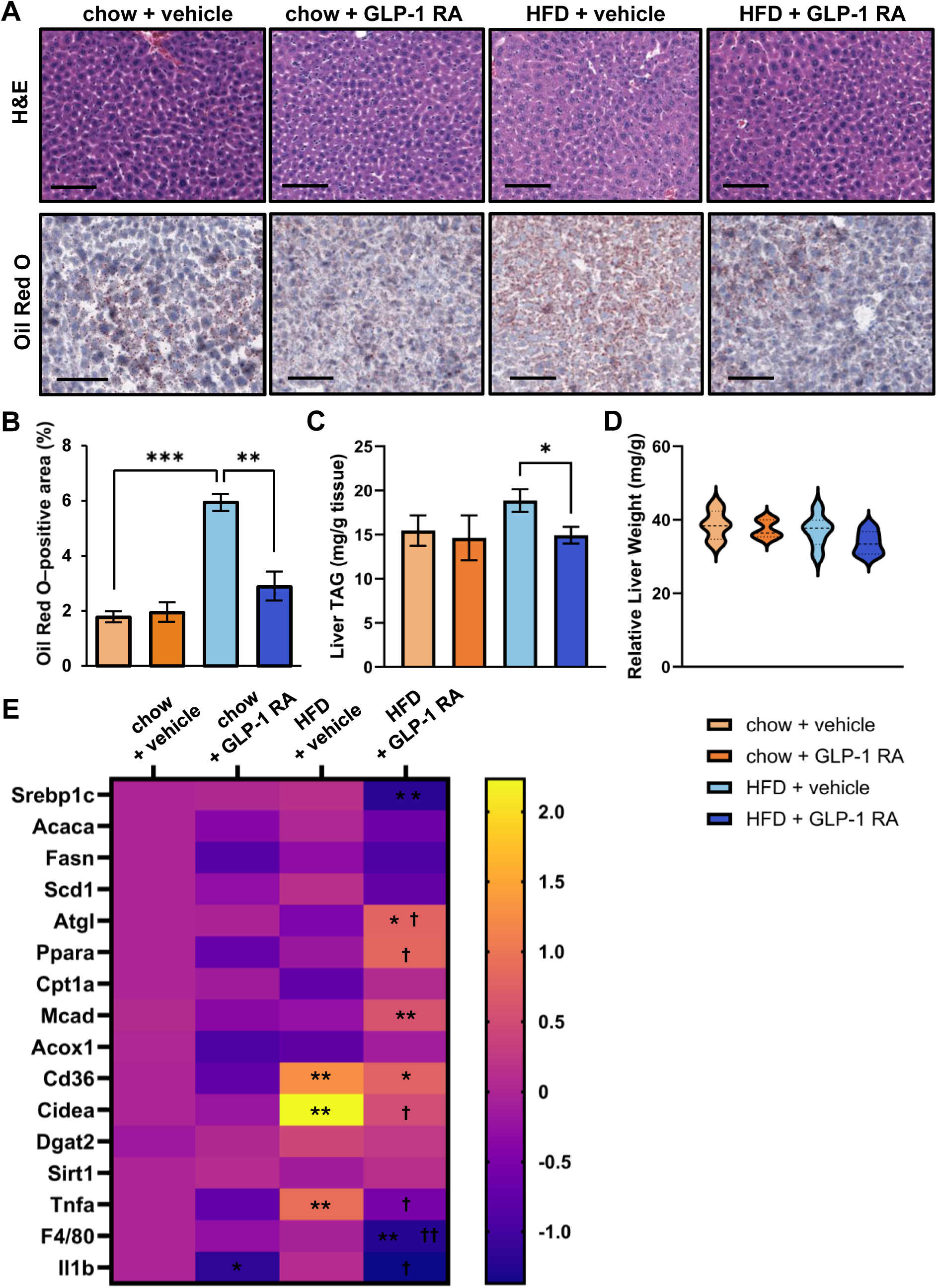
Hepatic lipid accumulation and gene expression in male F1 offspring at 18 weeks of age. (A) Representative hematoxylin and eosin (H&E) and Oil Red O (ORO) staining of liver sections from male F1 offspring at 18 weeks of age. Scale bar, 100 μm. (B) Quantification of ORO-positive area. (C) Hepatic triglyceride (TAG) content. (D) Relative liver weight. (E) Hepatic gene expression profiling of lipogenic, oxidative, lipid storage and transport, and inflammatory markers assessed by bulk quantitative PCR (qPCR), presented as log2 fold change in a heatmap. Data are presented as mean ± SEM (*n* = 6–9 male offspring). *P ≤ 0.05, **P ≤ 0.01, ***P ≤ 0.001 vs chow + vehicle. ^†^P ≤ 0.05, ^††^P ≤ 0.01 vs HFD + vehicle.

## DISCUSSION

In this study, maternal semaglutide treatment administered prior to conception and continued throughout gestation and lactation at a reduced dose improved maternal metabolic health during pregnancy and postpartum and attenuated long-term metabolic dysfunction in offspring, including reduced adiposity and hepatic steatosis and improved glucose homeostasis in adulthood. These findings demonstrate that pharmacologic optimization of maternal metabolic status prior to and during pregnancy can modulate intergenerational transmission of metabolic disease risk in a translationally relevant model.

Metabolic dysfunction during pregnancy, characterized by pre-pregnancy obesity, glucose dysregulation, and associated insulin resistance among women of reproductive age, is increasingly prevalent worldwide (35). Higher gestational glucose is associated with increased maternal risk of prediabetes and Type 2 diabetes years after pregnancy (19). Large multinational cohort studies including the Hyperglycemia and Adverse Pregnancy Outcomes (HAPO) study and subsequent follow-up studies which examined child outcomes at 10-14 years of age, have shown that subclinical metabolic dysfunction in pregnant women during the second trimester, characterized by elevated plasma glucose not diagnostic of gestational diabetes at that time, are associated with insulin resistance, impaired glucose metabolism, and increased adiposity in children independent of maternal BMI (18–20). Finally, preclinical studies have shown that a single generation of high-fat diet exposure promotes metabolic dysfunction in subsequent generations, with normalization requiring multiple generations of standard diet feeding (36–39). These data suggest that maternal metabolic dysfunction in utero creates durable impacts on both maternal and child health outcomes and presents an opportunity for prevention of metabolic disease that may extend across generations (11, 12, 16, 36).

Meta-analyses have shown that lifestyle modifications during pregnancy, including diet- and physical activity–based interventions, generally produce modest reductions in gestational weight gain and select outcomes such as cesarean delivery and gestational diabetes, without consistent improvements in broader adverse maternal or offspring composite outcomes (40, 41). Contemporary pharmacologic strategies in pregnancy focus on glycemic control with agents of established fetal safety, predominantly insulin; however, these approaches target overt hyperglycemia and do not address subclinical metabolic dysfunction that contributes substantially to future maternal diabetes risk and offspring obesity and diabetes (42, 43). In this context, GLP-1 RAs have emerged as highly effective agents to improve metabolic homeostasis in nonpregnant adults through both weight loss–dependent and –independent mechanisms, raising the question of whether preconception or periconception use could alter maternal metabolic risk trajectories and offspring outcomes.

While no clinical studies have directly evaluated GLP-1 receptor agonist use during gestation, emerging retrospective clinical studies have reported no increase in rates of congenital or cardiac malformations in children exposed in utero compared with other glucose-lowering therapies (44–46). GLP-1 RAs are currently not recommended for use during pregnancy due to limited clinical safety data, and existing clinical guidelines advise discontinuation before conception or as soon as pregnancy is recognized. However, a recent retrospective cohort study found that GLP-1 RA use with subsequent preconception or early pregnancy discontinuation was associated with greater gestational weight gain and higher risks of preterm delivery, gestational diabetes, and hypertensive disorders of pregnancy (47). These studies underscore the urgent need for rigorous evaluation of the safety and efficacy of GLP-1 RAs in the periconception and gestational periods, given their widespread use in women of reproductive age and their potential to alter maternal metabolic trajectories and intergenerational risk of obesity and diabetes.

Our lab previously investigated the therapeutic potential of exendin-4, a first-generation exogenous GLP-1 analog, in a rat model of uteroplacental insufficiency which predisposes offspring to diabetes (29, 30). A few preclinical studies performed in mouse models have investigated the safety and therapeutic potential of GLP-1 RAs during gestation (31, 48–52). However, many of these studies utilized supratherapeutic drug dosing with experimental designs not immediately translationally relevant, for example, with initiation of GLP- 1 RA treatment during gestation as opposed to pre-pregnancy or treatment only in chow-fed dams rather than in models of maternal obesity (31, 48, 49, 52). Critically, most preclinical studies assessing gestational GLP-1 RA exposure have not evaluated long-term outcomes in offspring, with some reports suggesting adverse short-term outcomes including reduced conception rate, altered placental structure, reduced fetal viability and skeletal abnormalities (31, 51). These findings highlight substantial heterogeneity in preclinical study design and a persistent gap in understanding the safety and long-term efficacy of clinically relevant GLP-1 receptor agonist exposure during pregnancy and lactation.

To our knowledge, this is the first preclinical study evaluating the safety and therapeutic potential of clinically relevant doses of semaglutide initiated prior to conception and continued throughout gestation and lactation in a diet-induced model of maternal obesity with assessment of long-term maternal and offspring metabolic outcomes (50). We show that preconception semaglutide promoted moderate weight loss of 5% over the course of 5 days. During gestation and lactation, semaglutide was administered at one-third of the preconception dose, a level at which no significant differences in body weight change was observed. At E17.5, a trend for reduction in fat mass was observed in HFD-fed dams treated with semaglutide, with a significant decrease in relative gonadal fat mass and adipocyte surface area at the microscopic level, suggesting overall reduction in adiposity and possible improvement in lipolytic and oxidative activity. Similar effects have been reported in non-pregnant preclinical models, where GLP-1 receptor agonism enhances lipolytic and oxidative activity in white adipose tissue, reduces adipose inflammation, and promotes thermogenic and browning gene expression, in some studies, independent of nutrient intake (53–55). We similarly observed significant improvements in glucose tolerance in HFD-fed dams treated with semaglutide at E15.5, despite the absence of weight loss during gestation. This is consistent with clinical data showing that liraglutide rapidly improves glycemic control and insulin sensitivity within 2 weeks of treatment in non-pregnant individuals with obesity and prediabetes prior to weight loss (56). We hypothesize that improvements in adiposity and glucose homeostasis during gestation between E14.5 through E17.5 is both a reflection of pre-conception weight loss and sustained improvements in white adipose tissue lipolytic activity and glucose-stimulated pancreatic β-cell insulin secretion during gestation with low-dose treatment (57).

Maternal semaglutide during pregnancy also reduced hepatic steatosis in HFD-fed animals. In parallel we saw marked changes in liver transcripts, most notably an increase in lipid droplet-associated and pro-inflammatory markers with HFD that were reduced with treatment. Indeed, several studies in non-pregnancy preclinical models have shown an improvement in steatosis with GLP-1 RA (57–62). Although most studies suggest minimal or absent GLP-1 receptor expression in hepatocytes, more recent work identifies GLP-1 receptor in other liver cell types, including liver endothelial cells, infiltrating monocytes and macrophages, and hepatic stellate cells (63, 64). In our diet-induced maternal obesity model, reductions in adiposity likely contributed substantially to the observed hepatic effects in semaglutide-treated mice. However, GLP-1 receptor agonism may also exert direct or indirect effects on hepatic non-parenchymal cell populations, as well as systemic inflammatory pathways independent of weight loss.

The relative contribution of weight loss–dependent and –independent mechanisms to these hepatic effects during pregnancy remains to be established.

Pregnancy reflects a state of physiological insulin resistance, and studies suggest that steatosis in settings of chronic overnutrition during gestation may worsen insulin sensitivity and contribute to the development of gestational diabetes, in addition to reduced pancreatic β-cell reserve pre-conception (65–68). In our model, maternal semaglutide-mediated reduction in hepatic steatosis suggests a potential mechanism for improved insulin sensitivity and glucose homeostasis, but follow-up studies specifically examining hepatic insulin sensitivity and pancreatic β-cell function are needed to better delineate the individual contributions of these pathways to the overall improvement in maternal metabolic health (59).

Our findings demonstrate that semaglutide’s metabolic benefits extend beyond the setting of diet-induced obesity, with beneficial effects observed in chow-fed dams during gestation. Notably, semaglutide reduced hepatic triglyceride content in chow-fed dams despite absence of differences in adiposity suggesting that GLP-1 receptor agonism may directly improve hepatic lipid handling independent of systemic fat mass—a finding consistent with non-pregnant preclinical studies showing weight loss–independent effects of GLP-1 RAs on liver metabolism (56, 61, 63). The trend toward improved glucose tolerance and increased lean mass in chow-fed semaglutide-treated animals further supports the notion that GLP-1 RAs may exert protective metabolic effects even in the absence of overt metabolic disease. Importantly, semaglutide exposure was not associated with adverse effects on conception rate, fetal viability, or fetal weight in chow- or HFD-fed animals under the dosing paradigm utilized in this study. These findings contrast with prior reports of adverse outcomes in chow-fed mice (31, 50, 51) and support further investigation into the safety and metabolic effects of clinically relevant GLP-1 RA exposure during pregnancy.

We next examined whether improvements in maternal metabolic health translated into long-term metabolic benefits in offspring maintained on a chow diet after weaning. Maternal semaglutide exposure in chow-fed dams had minimal effects on long-term offspring metabolic phenotypes. However, offspring exposed to maternal HFD displayed evidence of persistent metabolic dysfunction despite postnatal chow feeding, including accelerated early postnatal weight gain, increased adiposity, impaired glucose homeostasis, and hepatic steatosis in adulthood. Importantly, these adverse metabolic effects were mitigated by maternal semaglutide treatment.

The beneficial effects of maternal semaglutide exposure in HFD-exposed male offspring were particularly evident at the tissue level, including improvements in adipose tissue remodeling and hepatic metabolism. Maternal HFD exposure promoted adipocyte hypertrophy and altered adipocyte size distribution in adult male offspring, consistent with prior studies linking maternal obesity to long-term adipose tissue dysfunction and increased metabolic disease risk in offspring (69, 70). These changes were accompanied by hyperinsulinemia, increased hepatic triglyceride accumulation, and increased expression of hepatic lipid storage and inflammatory transcripts, which were ameliorated by maternal semaglutide exposure. Offspring from HFD-fed dams exposed to semaglutide also demonstrated reduced expression of lipogenic genes and increased expression of oxidative markers, suggesting persistent alterations in hepatic metabolic programming. While several developmental programming models suggest hepatic metabolic adaptations (5, 71, 72), the improvements in glucose homeostasis and hepatic triglyceride content observed in offspring from semaglutide-treated HFD-fed dams likely reflect both systemic effects related to reduced adiposity and direct effects on tissue-specific metabolic programming. Further studies are required to delineate the relative contributions of systemic versus tissue-specific mechanisms of developmental programming in this model.

Sex-specific differences in developmental programming have been widely reported (69, 73, 74), with some studies showing male offspring demonstrating greater susceptibility to metabolic dysfunction compared with females (75). Consistent with this, the metabolic phenotype in female offspring exposed to maternal HFD in the present study was comparatively attenuated, with no significant differences in body weight, adiposity, or glucose tolerance compared with controls and only a modest impairment in insulin tolerance observed. Maternal semaglutide did not significantly alter metabolic outcomes in female offspring.

Our experimental design did not distinguish between the relative contributions of gestational versus lactational semaglutide exposure on offspring outcomes. GLP-1 receptor expression in the placenta remains controversial, and it is unclear whether semaglutide crosses the placenta to directly influence fetal tissues (50, 76, 77).

Similarly, transfer into breast milk and potential effects on postnatal development have not been fully established (78). We hypothesize that the long-term benefits observed in offspring exposed to maternal semaglutide primarily reflect improvements in maternal metabolic status rather than direct drug exposure, although future studies measuring drug levels in fetal and postnatal tissues, along with temporal separation of gestational versus lactational exposure, are required to distinguish these mechanisms.

In addition to assessing long-term outcomes in offspring, we evaluated postpartum outcomes in dams (F0) following offspring weaning and treatment discontinuation. HFD-fed dams previously exposed to semaglutide had reduced adiposity, without changes in lean mass or bone mineral density, at 10 weeks postpartum. Most notably, prior treatment with semaglutide promoted persistent improvements in glycemic control and insulin tolerance in HFD-fed primiparous mice 4 and 8 weeks after treatment discontinuation, respectively. It is possible that early improvements in metabolic status prior to conception, together with continued semaglutide exposure during gestation and lactation, contributed to the sustained improvements in adiposity and glucose homeostasis observed after treatment discontinuation. However, clinical data in non-pregnant individuals consistently show that discontinuation of GLP-1 RA results in weight rebound and worsening glycemic parameters (79). Our data suggests that pregnancy represents a critical window during which alterations in metabolic homeostasis may promote durable changes in maternal metabolism. Indeed, recent data demonstrate that pregnancy can reprogram maternal energy metabolism and induce long-term postpartum obesity via altered adipose tissue signaling and reduced energy expenditure, independent of fat retention during pregnancy (80). At the cellular level, maternal hepatocytes undergo heterogeneous and dynamic developmental phenotypic changes during gestation indicating that pregnancy drives cell-type-specific metabolic reprogramming in maternal organs (81). Clinical studies further support pregnancy as a critical window for long-term metabolic risk, with excessive gestational weight gain and metabolic complications associated with persistent alterations in maternal body composition, insulin resistance, and cardiovascular risk years later (82, 83). However, strategies initiated prior to conception and maintained throughout pregnancy to improve maternal metabolic health and prevent adverse metabolic programming in both dams and offspring remain underinvestigated. In this preclinical study, we address this gap using a model of maternal obesity and GLP-1 RA exposure. Our findings provide evidence that targeting maternal metabolic dysfunction during critical developmental windows may confer benefits to both maternal and offspring long-term metabolic health.

Metabolic dysfunction represents a rapidly growing global health crisis, affecting nearly 1.5 billion individuals worldwide (84). The continued rise in obesity and metabolic disease, despite the development of highly effective anti-obesity therapeutics, may be driven in part by the intergenerational transmission of metabolic risk (11, 12, 16). Using a diet-induced mouse model of maternal obesity and metabolic dysfunction, we demonstrate that preconception, gestational, and lactational semaglutide exposure improves maternal metabolic health during pregnancy and postpartum and mitigates adiposity, glycemic dysregulation, and hepatic steatosis in adult offspring. These findings support the need for further studies evaluating the safety, efficacy, and mechanisms of GLP-1 RAs and related biologics as therapeutic strategies to treat maternal metabolic dysfunction and potentially prevent the developmental programming of metabolic disease in offspring. Collectively, these data provide a foundation for future mechanistic and translational studies investigating interventions targeting maternal metabolism during critical developmental windows.

## ACKNOWLEDGEMENTS

The authors thank the Center for Molecular Studies in Digestive and Liver Diseases (P30DK050306) and the Molecular Pathology and Imaging Core (RRID: SCR_022420) for technical support. The authors also acknowledge the University of Pennsylvania Rodent Metabolic Phenotyping Core for assistance with metabolic phenotyping, including EchoMRI and DEXA PIXImus analyses.

## FUNDING

This work was supported by Novo Nordisk, which provided semaglutide and contributed to discussions regarding dosing strategy and study design. The funder had no role in data collection, data analysis, data interpretation, or manuscript preparation.

A.R. was supported in part by the NIH T32 Clinical Pharmacology Training Program (T32HD117733-01).

## CONFLICTS OF INTEREST

The authors declare no competing interests.

### AUTHOR CONTRIBUTIONS

R.A.S. conceived the study. A.R. contributed to study design and performed experiments, data analysis, and manuscript preparation. T.H. and Y.G. assisted with animal studies and data collection. S.E.P. and R.A.S. provided critical revisions and supervised the project. All authors reviewed and approved the final manuscript.

## Notes

### Competing Interest Statement

The authors have declared no competing interest.

