## Supplemental Materials for "Perinatal Semaglutide Treatment Improves Maternal Health and Mitigates Offspring Metabolic Dysfunction in a Mouse Model of Maternal Obesity"

**Table S1. Mouse qPCR primer sequences for hepatic gene expression analysis.**

| Gene | Primer | Sequence (5'→3') |
|---|---|---|
| 36b4 | Forward | GGTGCCTCTGGAGATTTTCG |
| 36b4 | Reverse | CACTGGTCTAGGACCCGAGAAG |
| Srebp1c | Forward | GGAGCCATGGATTGCACATT |
| Srebp1c | Reverse | GGCCCGGGAAGTCACTGT |
| Acaca | Forward | GACAGACTGATCGCAGAGAAAG |
| Acaca | Reverse | TGGAGAGCCCCACACACA |
| Fasn | Forward | GCTGCGGAAACTTCAGGAAAT |
| Fasn | Reverse | AGAGACGTGTCACTCCTGGACTT |
| Scd1 | Forward | CCGGAGACCCCTTAGATCGA |
| Scd1 | Reverse | TAGCCTGTAAAAGATTTCTGCAAACC |
| Atgl | Forward | GCCTCCTTGGACACCTCAATAA |
| Atgl | Reverse | CTTCCTCGGGGTCTACCACA |
| Ppara | Forward | CACCTGCAGAGCAACCATC |
| Ppara | Reverse | CCGAAGGTCCACCATTTTT |
| Cpt1a | Forward | TGAGTGGCGTCCTCTTTGG |
| Cpt1a | Reverse | CAGCGAGTAGCGCATAGTCATG |
| Mcad | Forward | TTACCGAAGAGTTGGCGTATG |
| Mcad | Reverse | ATCTTCTGGCCGTTGATAACA |
| Acox1 | Forward | CAGCAGGAGAAATGGATGCA |
| Acox1 | Reverse | GGGCGTAGGTGCCAATTATCT |
| Cd36 | Forward | TGTGTTTGGAGGCATTCTCA |
| Cd36 | Reverse | TTTTGCACGTCAAAGATCCA |
| Cidea | Forward | CTCCGAGTACTGGGCGATAC |
| Cidea | Reverse | ACCAGCCTTTGGTGCTAGG |
| Fsp27 | Forward | GGCTCACAGCTTGGAGGA |
| Fsp27 | Reverse | CTCCACGATTGTGCCATCT |
| Plin2 | Forward | GCTGCGGAAACTTCAGGAAAT |
| Plin2 | Reverse | AGAGACGTGTCACTCCTGGACTT |
| Dgat2 | Forward | AGCTGGTGAAGACACACAACC |
| Dgat2 | Reverse | TGATGATAGCATTGCCACTCC |
| Sirt1 | Forward | ATCGGCTACCGAGACAAC |
| Sirt1 | Reverse | GTCACTAGAGCTGGCGTGT |
| Tnfa | Forward | AGCACAGAAAGCATGATCCG |
| Tnfa | Reverse | CCCGAAGTTCAGTAGACAGAAGAG |
| F4/80 | Forward | CTTTGGCTATGGGCTTCCAGTC |
| F4/80 | Reverse | GCAAGGAGGACAGAGTTTATCGTG |
| Il1b | Forward | GAATGACCTGTTCTTTGAAGTT |
| Il1b | Reverse | TTTTGTTGTTCATCTCGGAGCC |
| Timp1 | Forward | CCTTTGCATCTCTGGCATCT |
| Timp1 | Reverse | CTCGTTGATTTCTGGGGAAC |

**Supplementary Figure 1. Maternal food intake during treatment and pregnancy phases.**

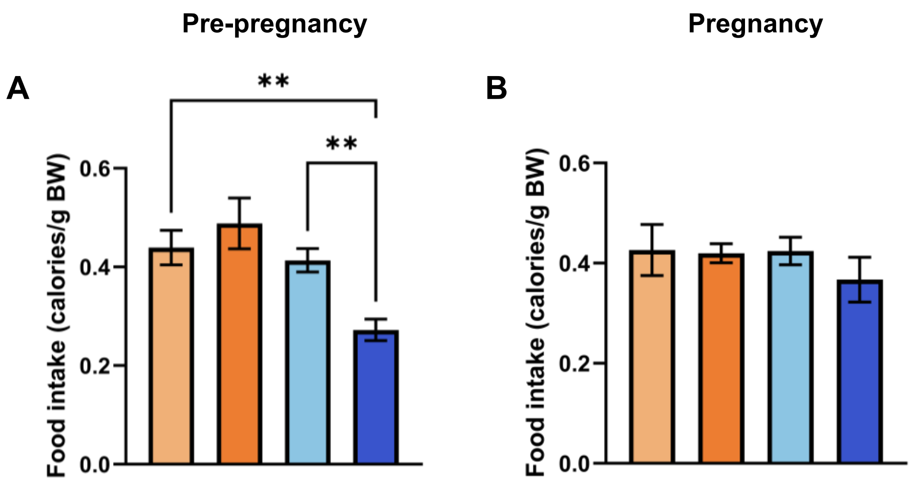


**Supplementary Figure 1. Maternal food intake during treatment and pregnancy phases.** (A) Food intake during the pre-pregnancy treatment phase (day 5). (B) Food intake during pregnancy at embryonic day (E)10.5. Data are presented as mean ± SEM (*n* = 10–12 chow, *n* = 15–16 HFD). **P ≤ 0.01.

**Supplementary Figure 2. Pregnancy outcomes and placental morphology at E17.5.**

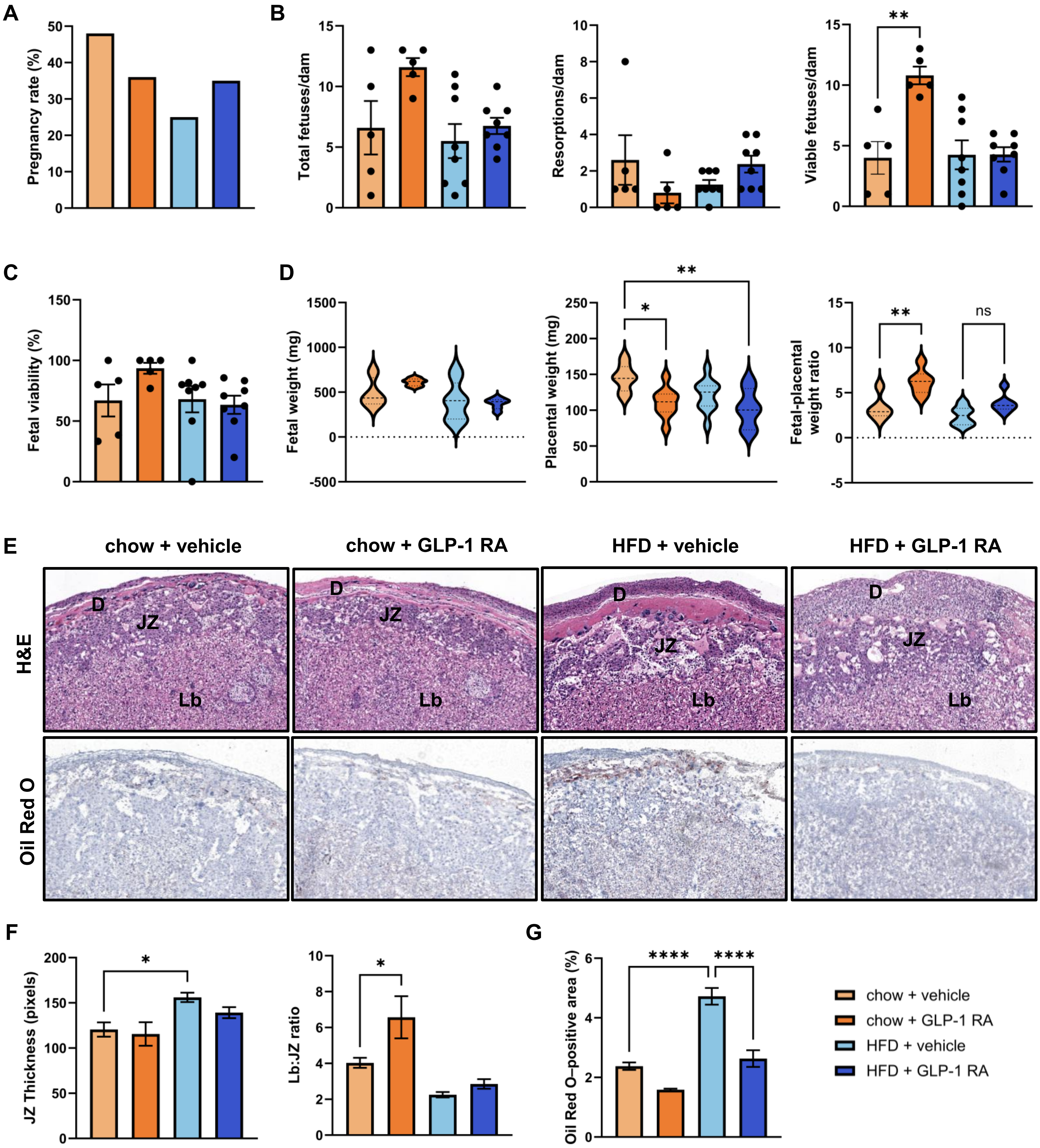

**Supplementary Figure 2. Pregnancy outcomes and placental morphology at E17.5.** (A) Pregnancy rate. (B) Total fetuses per dam, resorptions per dam, and viable fetuses per dam. (C) Fetal viability (%). (D) Fetal weight, placental weight, and fetal-to-placental weight ratio at embryonic day (E)17.5. (E) Representative hematoxylin and eosin (H&E, top) and Oil Red O (ORO, bottom) staining of placenta. (F) Junctional zone (JZ) thickness and labyrinth-to-junctional zone (Lb:JZ) ratio. (G) Placental ORO–positive area. Lb, labyrinth; JZ, junctional zone. Data are presented as mean ± SEM (chow: $n$ = 5 litters; HFD: $n$ = 8 litters). Pregnancy rate was analyzed using Fisher's exact test; all other comparisons were analyzed using one-way ANOVA. *P ≤ 0.05, **P ≤ 0.01, ****P ≤ 0.0001.

**Supplementary Figure 3. Glucose and insulin tolerance in female F1 offspring.**

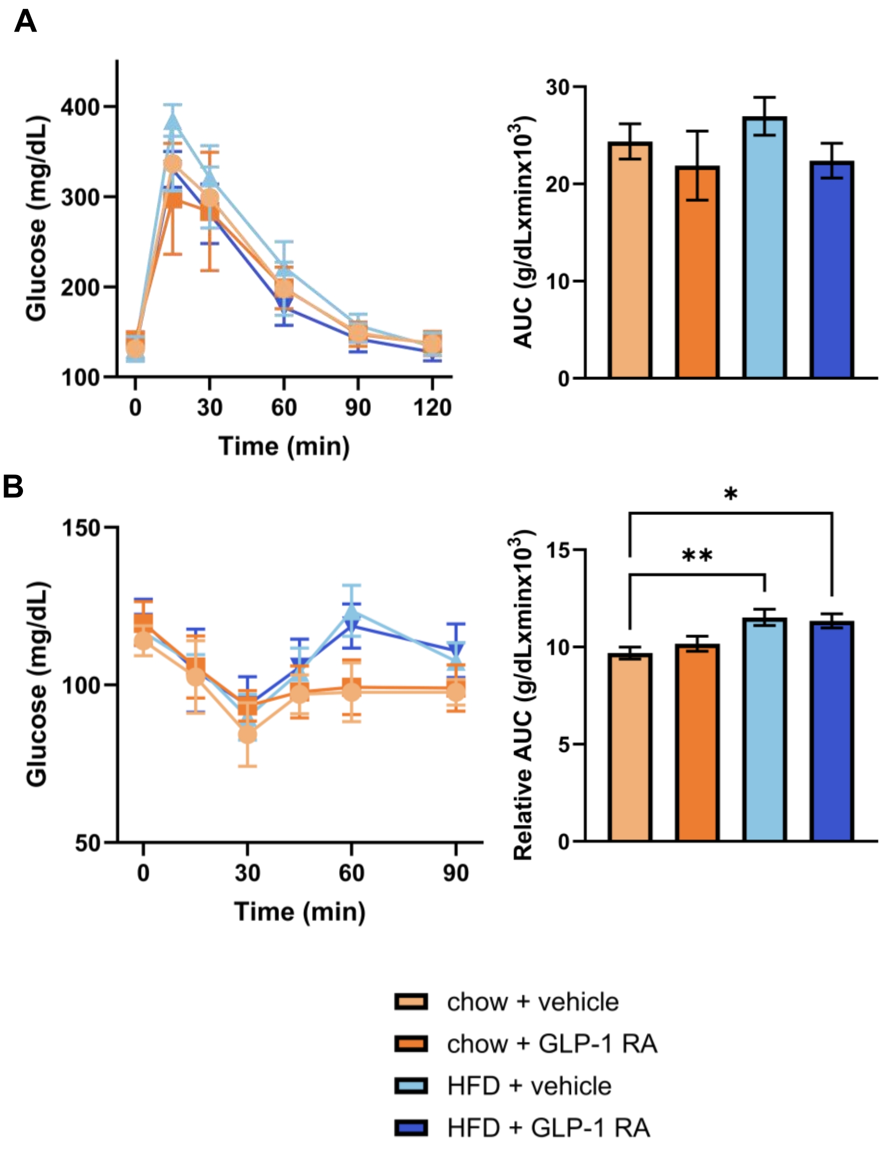


**Supplementary Figure 3. Glucose and insulin tolerance in female F1 offspring.** (A) Glucose tolerance test (GTT) at 8 weeks of age and insulin tolerance test (ITT) at 12 weeks of age in female F1 offspring. Area under the curve (AUC) for GTT and relative AUC for ITT are shown. Data are presented as mean ± SEM (*n* = 6–9 female offspring).
